## Supplementary Figures and Tables for "CEP peptide and cytokinin pathways converge on CEPD glutaredoxins to inhibit root growth"

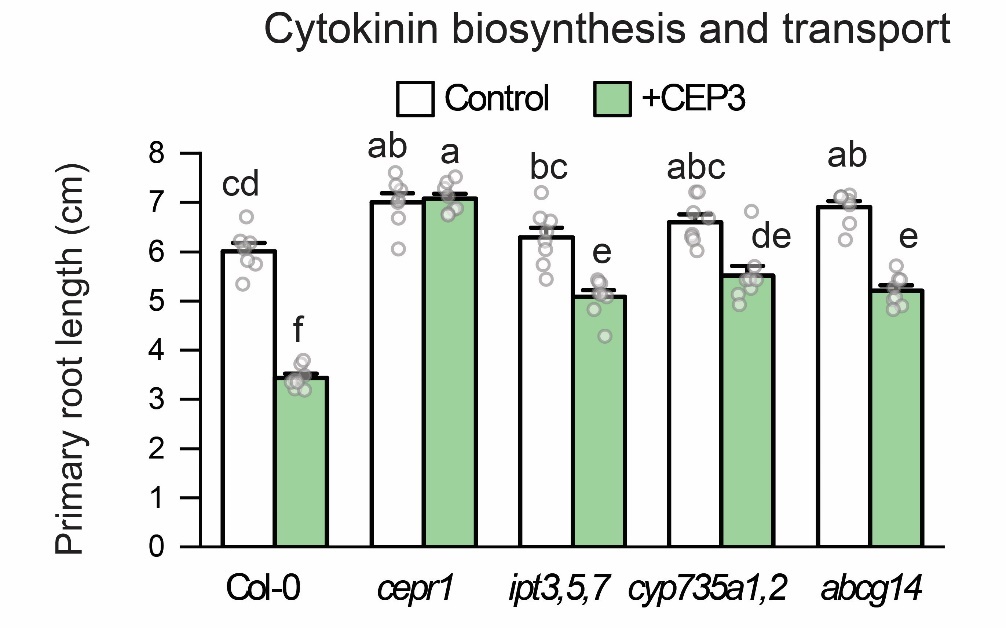

**Supplementary Figure 1. Absolute root growth measurements corresponding to Figure 1.** Primary root length for cytokinin biosynthesis or transport mutants grown on medium with or without CEP3 peptide (10^-6^ M) for 10 days in comparison to Col-0 and *cepr1-3* (n = 7-8 plants). Root length expressed as a percentage of seedlings grown on medium without CEP3 (control) for each respective genotype. Letters show significant differences (ANOVA followed by Tukey HSD test, p < 0.05).

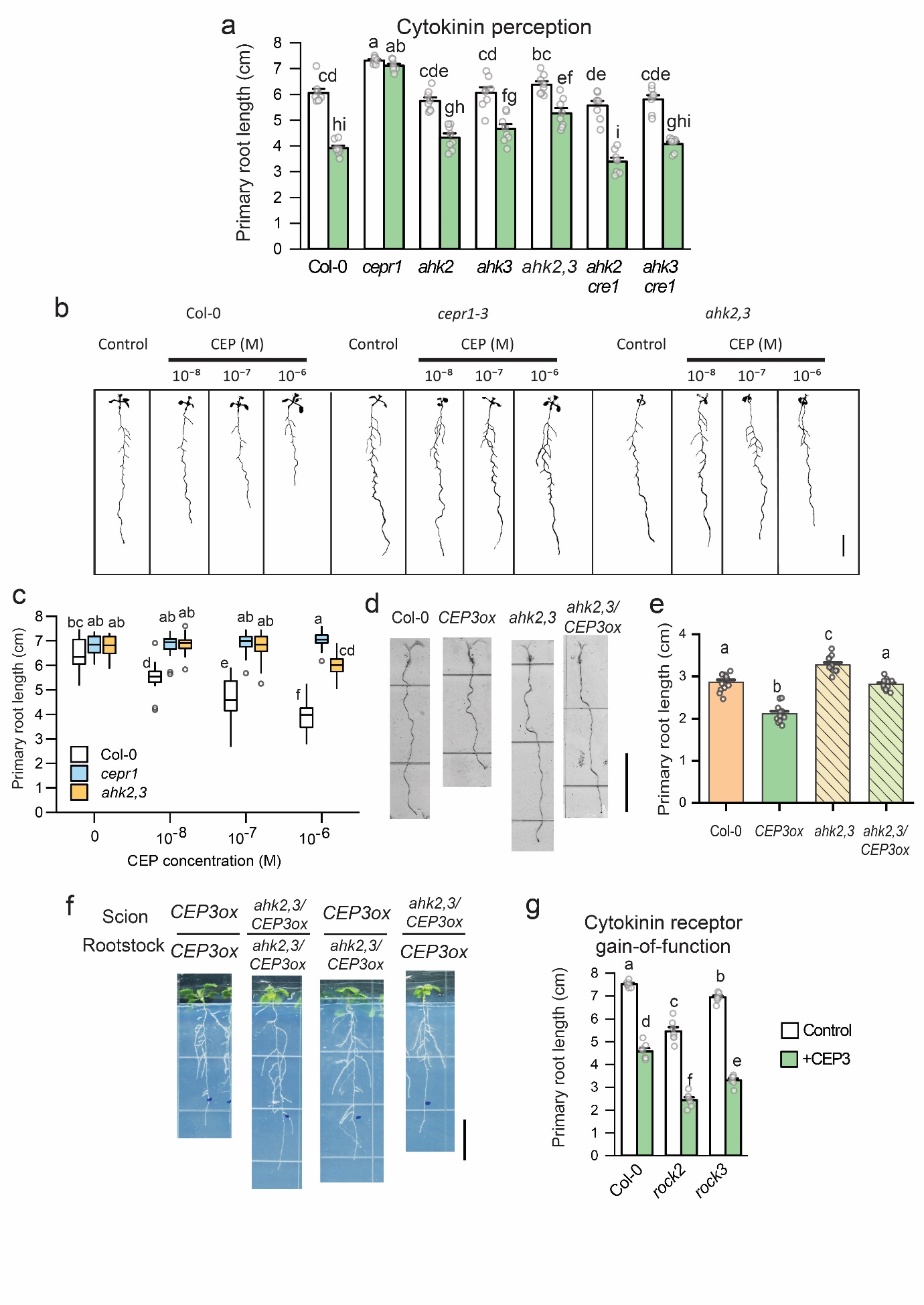

**Supplementary Figure 2. Absolute root growth measurements and representative images corresponding to Figure 2. a** Primary root length for cytokinin perception mutants grown on medium with or without CEP3 peptide (10^-6^ M) for 10 days in comparison to Col-0 and *cepr1-3* (n = 8 plants). **b** Representative images and **c** primary root length for Col-0, *cepr1* and *ahk2,3* in response to increasing CEP3 concentrations (0, 10^-8^, 10^-7^ or 10^-6^ M) at 10 days of growth (n = 17-21 plants). **d** Representative images and **e** primary root length of *CEP3* overexpressing plants in wild-type and *ahk2,3* backgrounds after 7 days growth (n = 12). **f** Representative images for reciprocal hypocotyl grafts between *CEP3* overexpressing plants in the Col-0 and *ahk2,3* backgrounds 12 days post grafting. Blue dots show primary root length 5 days post recovery from grafting **g** Primary root length of gain-of-function AHK mutants grown on medium with or without CEP3 peptide (10^-6^ M) for 12 days in comparison to Col-0. Letters indicate significant differences (ANOVA followed by Tukey HSD test, p < 0.05). Error bars show SE. Scale bars = 1 cm.

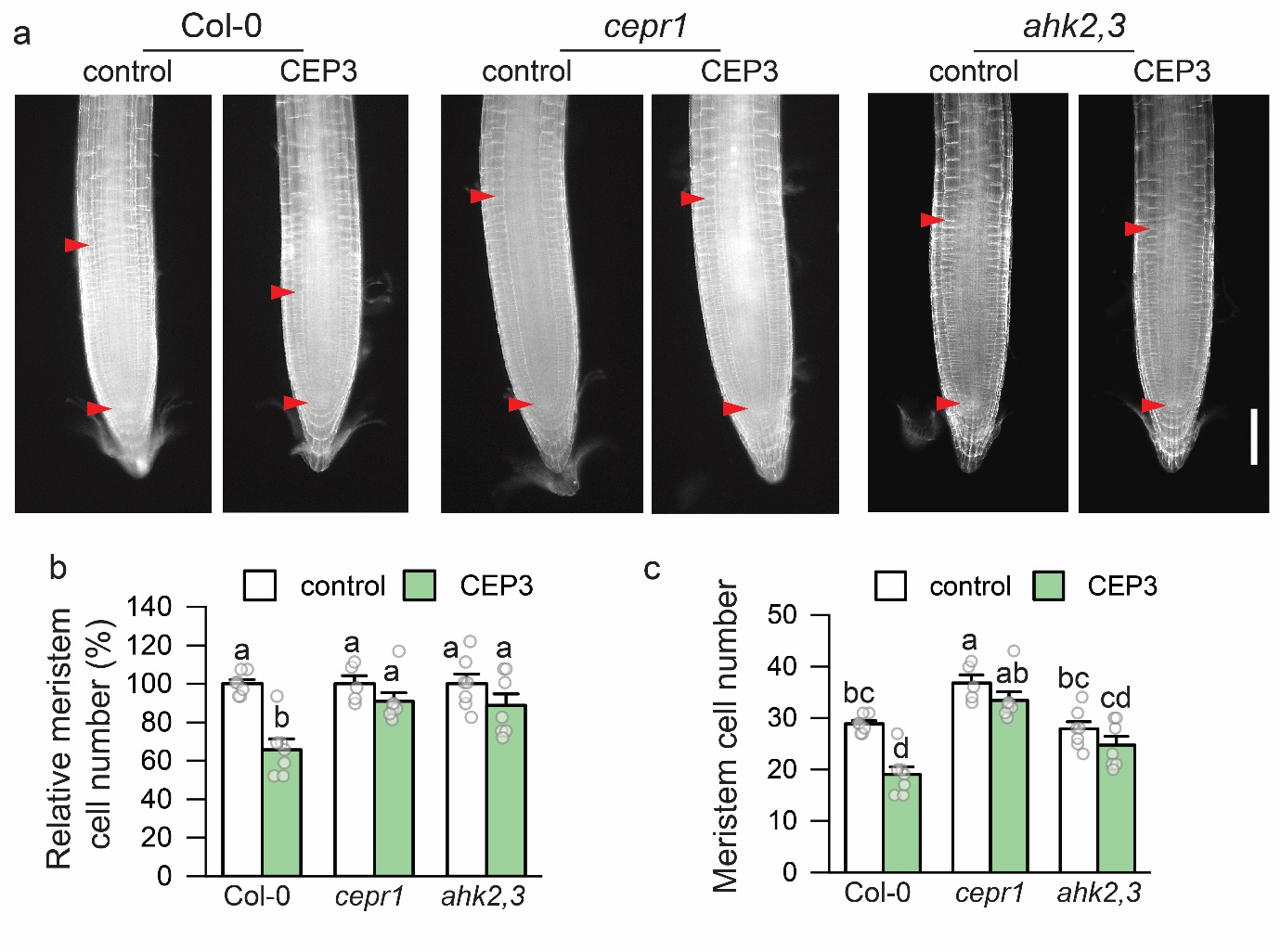

**Supplementary Figure 3. CEP inhibition of root apical meristem cell number depends on AHK2,3 activity.** Col-0, *cepr1-3* and *ahk2,3* plants were grown for 5 days before transfer to plates with or without CEP3 (1 µM) for 24 h. **a** Representative images of propidium iodide stained primary root tips, with the meristem boundaries indicated with red arrows. **b** Relative Meristem cortical cell number as a percentage of the respective control group for each line, and **c** absolute meristem cortical cell number. n = 5-7. Letters indicate significant differences (ANOVA followed by Tukey HSD test, p < 0.05). Error bars show SE. Scale bar = 100 µm.

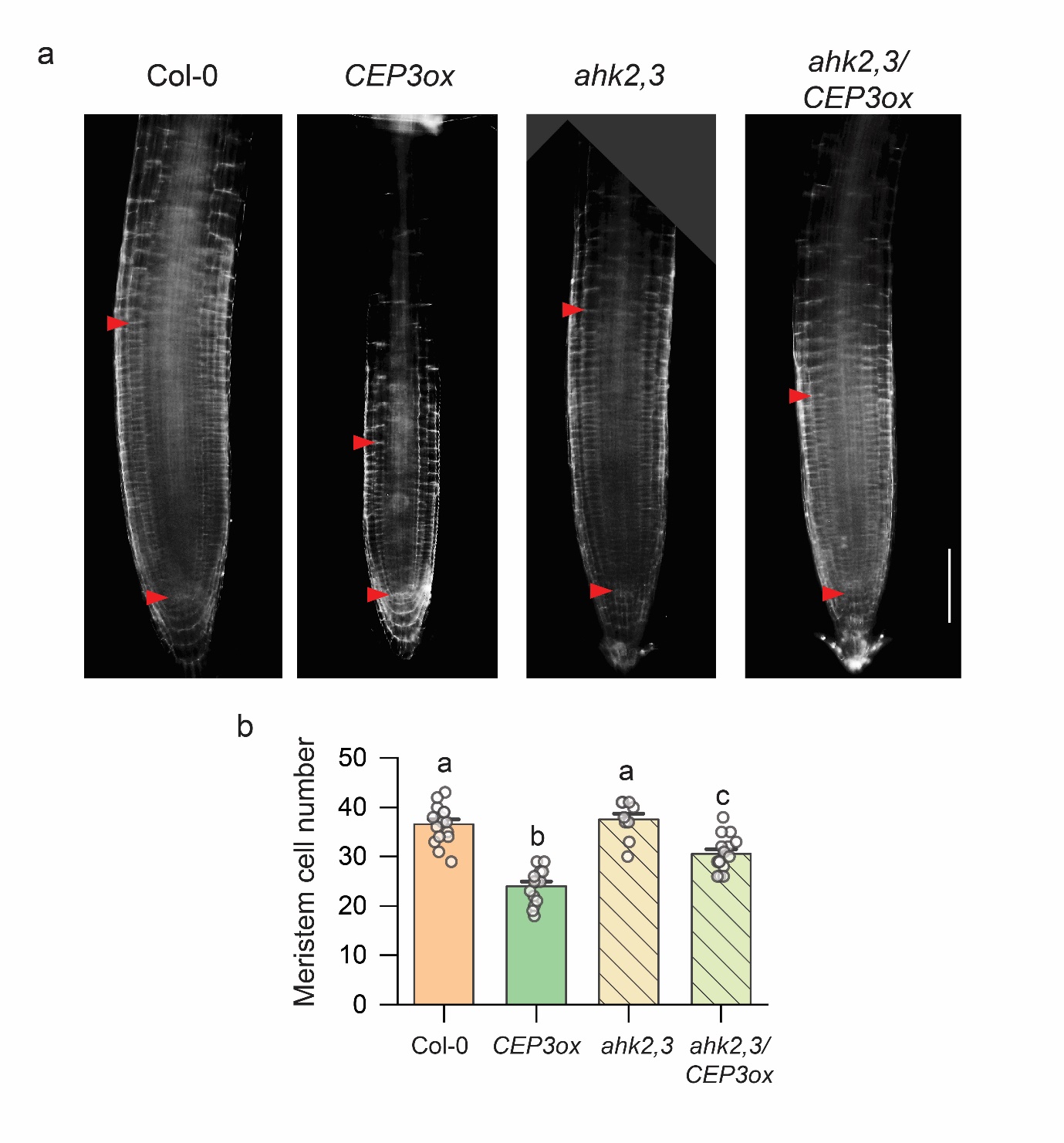

**Supplementary Figure 4. Inhibition of root apical meristem cell number by *CEP3* overexpression depends on AHK2,3 activity.** Col-0, *CEP3ox*, *ahk2,3*, and *ahk2,3/CEP3ox* plants were grown for 6 days. **a** Representative images of propidium iodide stained primary root tips, with the meristem boundaries indicated with red arrows. **b** Meristem cortical cell number. n = 10-16. Letters indicate significant differences (ANOVA followed by Tukey HSD test, p < 0.05). Error bars show SE. Scale bar in = 100 µm.

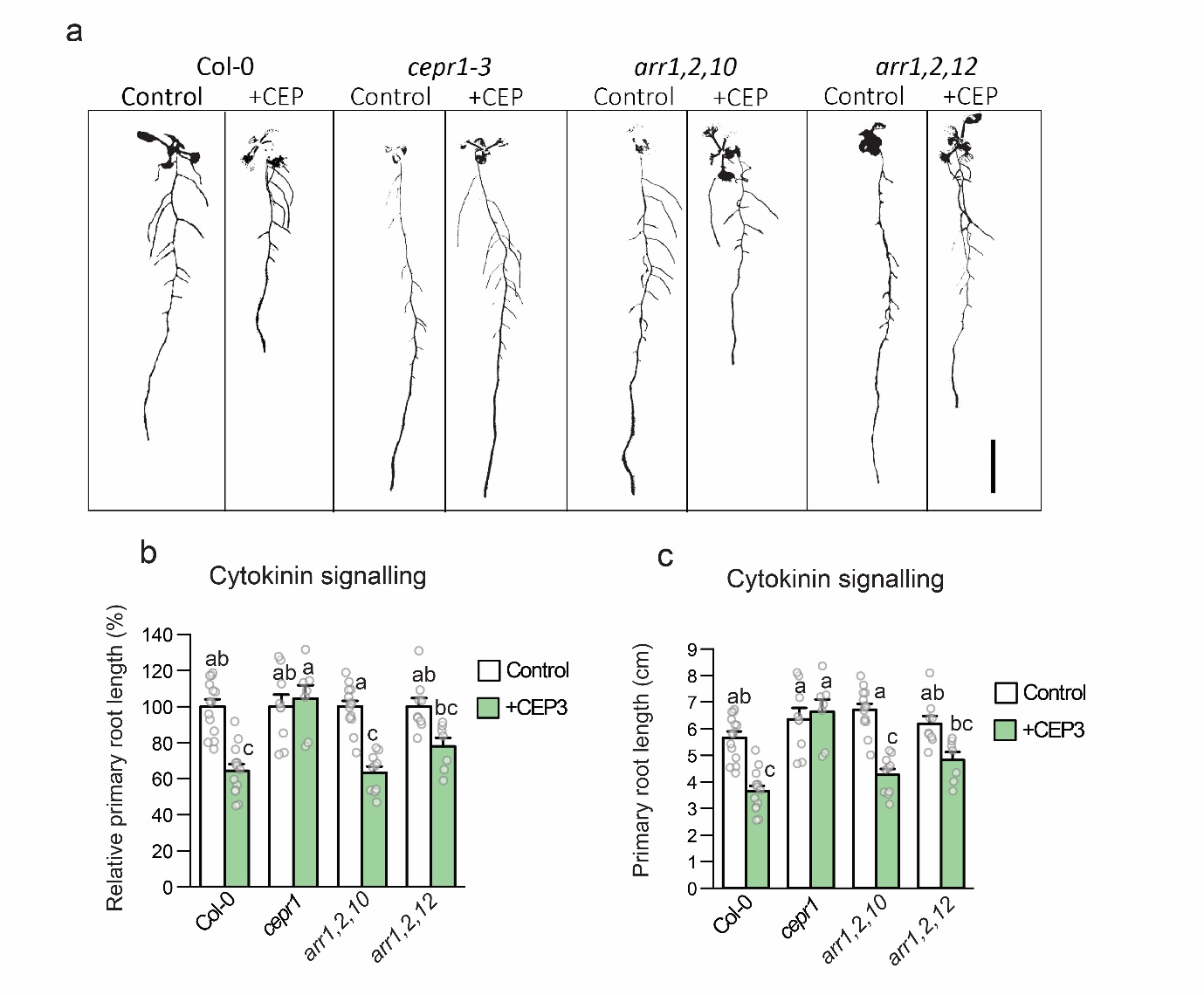

**Supplementary Figure 5. CEP sensitivity depends on cytokinin signalling via ARRs. a** Representative images, **b** relative primary root length, and **c** absolute primary root growth of Col-0, *cepr1-3* and mutants affected in cytokinin signalling. Plants were grown for 11 days on medium with or without CEP3 (10^-6^ M) (n = 7-14). Relative root length in **b** expressed as a percentage of seedlings grown on medium without CEP3 (control) for each respective genotype. Letters indicate significant differences (ANOVA followed by Tukey HSD test, p < 0.05). Error bars show SE. Scale bar = 1 cm.

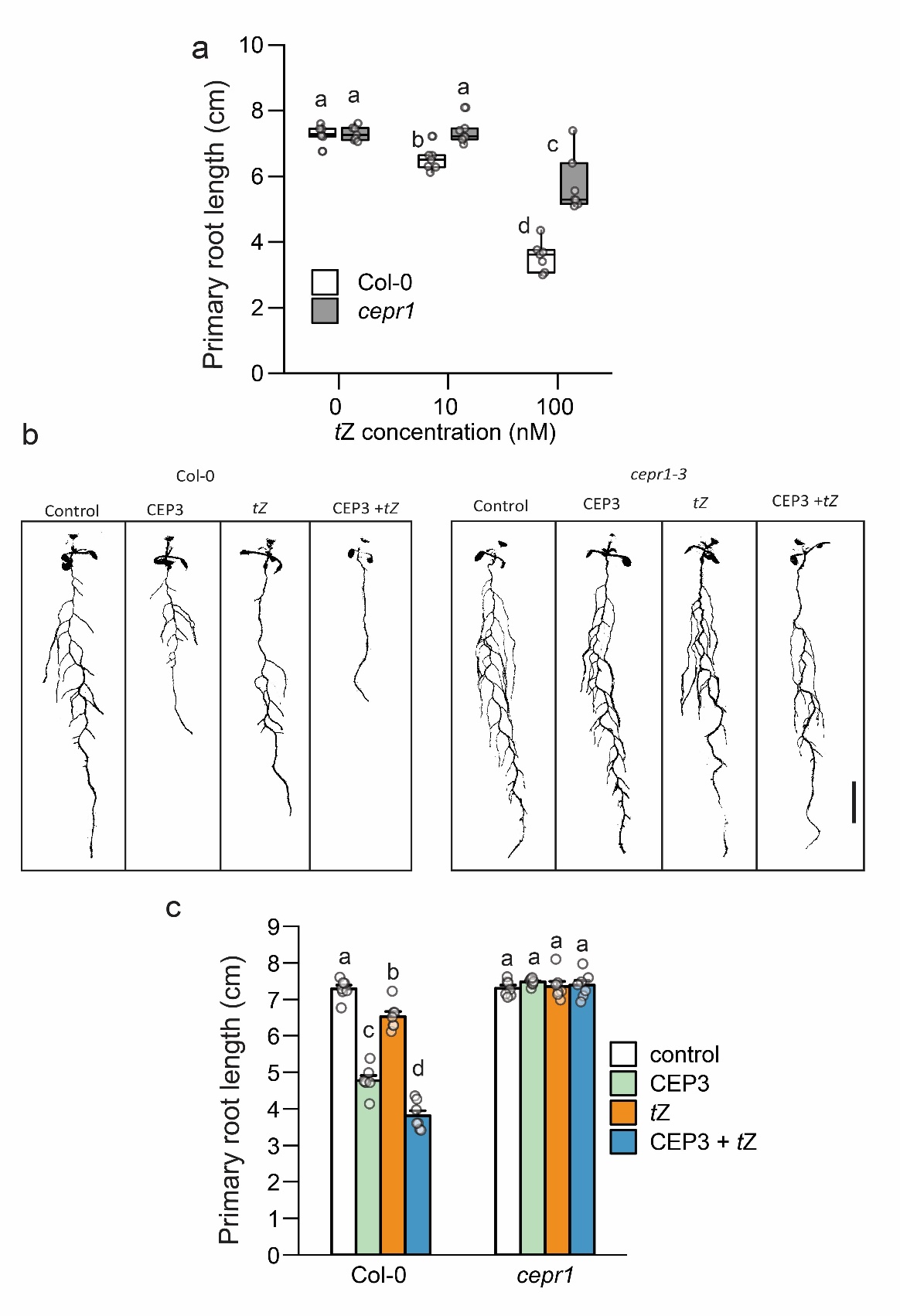

**Supplementary Figure 6. Absolute root growth measurements and representative images corresponding to Figure 3. a** Primary root length for Col-0 and *cepr1-3* in response to increasing concentrations of *t*Z (0, 10, 100 nM) at 12 dg (n=7). **b,c** CEP and *t*Z signalling is defective in *cepr1-3.* **b** Representative images and **c** primary root length for Col-0 and *cepr1-3* plants grown on medium containing CEP3 (1 µM), *t*Z (10 nM), combined CEP3 and *t*Z, or control medium (no CEP or tZ) for 12 days (n = 7) (ANOVA followed by Tukey HSD test, p < 0.05). Error bars show SE. Scale bar = 1 cm.

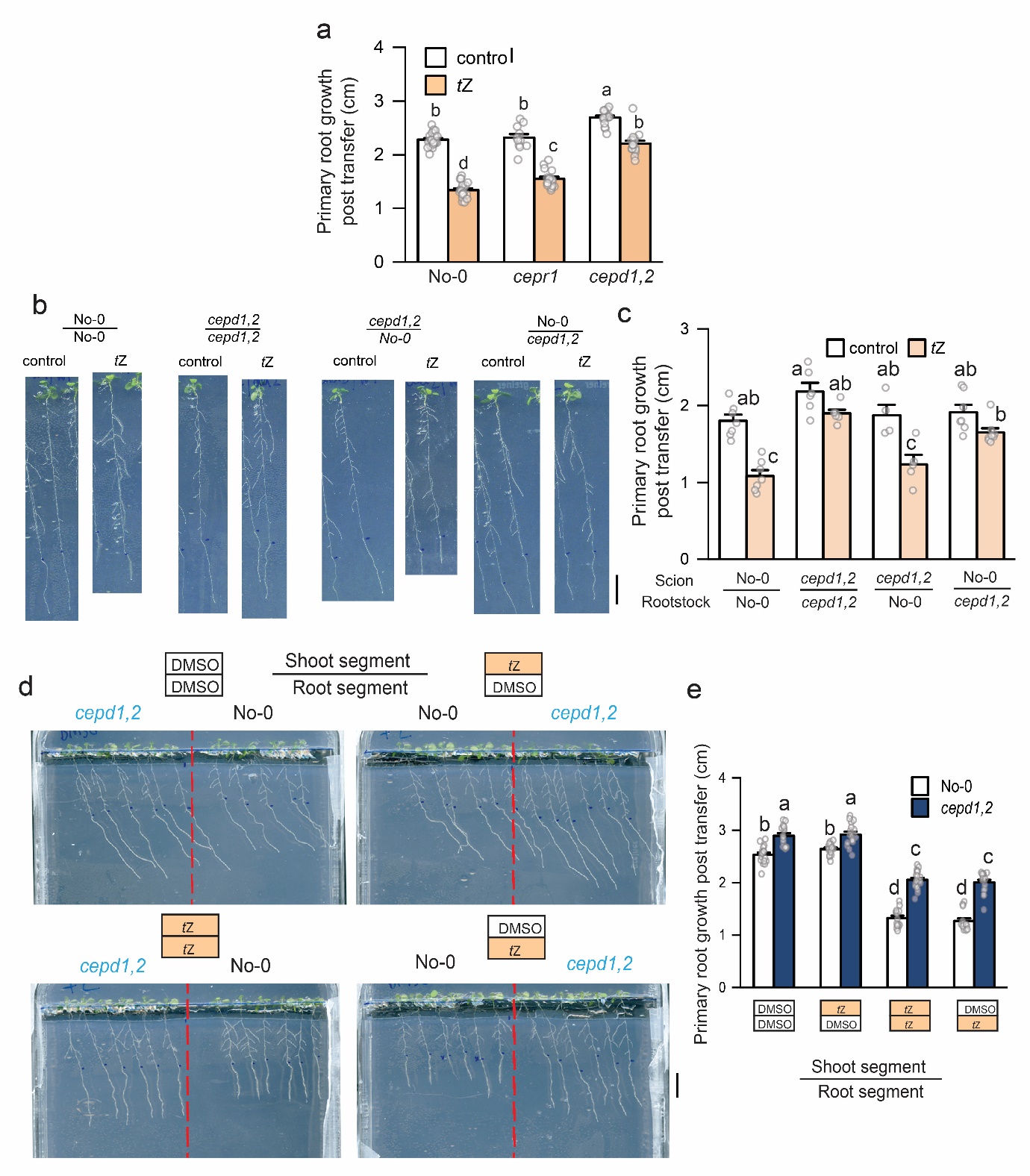

**Supplementary Figure 7. Absolute root growth measurements and representative images corresponding to Figure 5.** Primary root growth for No-0, *cepr1-1*, and the *cepd1,2* double mutant seedlings 3 days after transfer to medium with or without *t*Z (10 nM) (n = 11-24). **b,c** *t*Z sensitivity of reciprocal hypocotyl grafts between No-0 and *cepd1,2*. **b** Representative images and **c** primary root growth 3 days post transfer to *t*Z (5 nM) for each respective graft combination (n = 4-8). Blue dots in **b** indicate the length of the primary root on the day of transfer. **d** Representative images of a segmented agar plate set-up for *t*Z treatment of roots and/or shoots. Plants were grown for 6 days before transfer to segmented plates with tZ (10 nM) or solvent control (DMSO) infused in each plate segment. Blue dots indicate root tips on day of transfer. Plants were positioned such that roots only were in contact with the bottom segment, and shoots only were in contact with the top segment. **e** Primary root growth for No-0 and *cepd1,2* seedlings 3 days post transfer to segmented plates (n = 13-18). Root growth post transfer expressed as a percentage of the DMSO control for each respective genotype. Letters show significant differences (ANOVA followed by Tukey HSD test, p < 0.05). Error bars show SE. Scale bars = 1 cm.

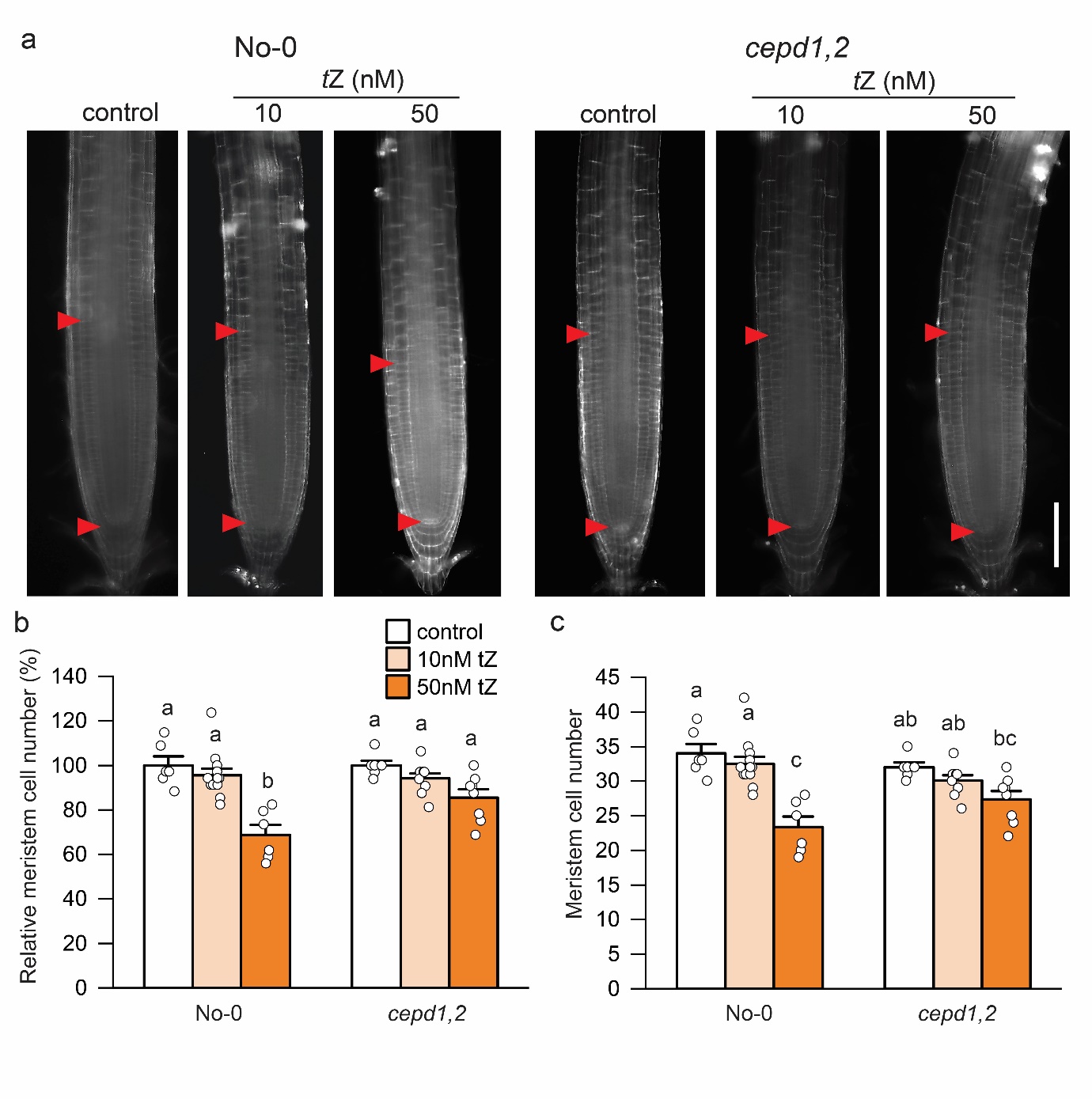
**Supplementary Figure 8. *t*Z inhibition of root apical meristem cell number depends on CEPD activity.** No-0 and *cepd1,2* plants were grown for 5 days before transfer to plates containing DMSO (control), 10nM *t*Z, or 50nM *t*Z. **a** Representative images of propidium iodide-stained primary root tips, with the meristem boundaries indicated with red arrows. **b** Relative Meristem cortical cell number as a percentage of the respective control group for each line, and **c** absolute meristem cortical cell number (n = 6-12). Letters indicate significant differences (ANOVA followed by Tukey HSD test, p < 0.05). Error bars show SE. Scale bar = 100 µm.

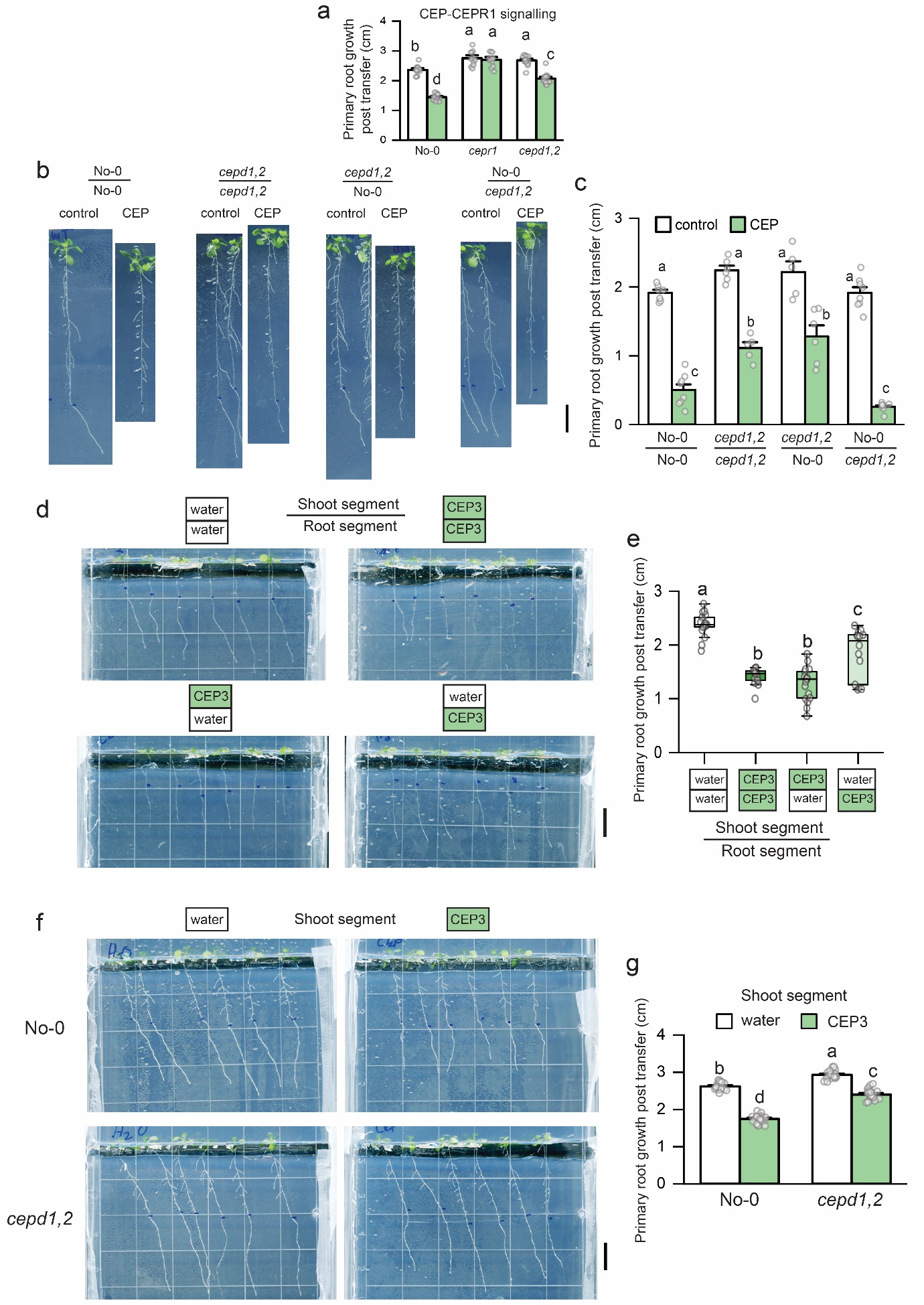

**Supplementary Figure 9. Absolute root growth measurements, representative images and extended data associated with Figure 6. a** Primary root growth for No-0, *cepr1-1*, and the *cepd1,2* double mutant seedlings 3 days after transfer to medium with or without CEP3 (10^-6^ M) (n=12-18). **b,c** CEP3 sensitivity of reciprocal hypocotyl grafts between No-0 and *cepd1,2*. **a** Representative images and **b** primary root growth 3 days post transfer to treatment plates (n = 5-8). **d** Representative images of a segmented agar plate set-up for selective CEP3 treatment of shoots and roots. Col-0 were grown for 6 days before transfer to segmented plates with CEP3 (10^-6^ M) or solvent control (water) infused in the top or bottom agar segments. Plants were positioned such that shoots but not roots were in contact with the top segment and plants were grown for a further 3 days after transfer. **e** Primary root growth post transfer to segmented plates (n = 14-17 plants). **f** Representative images and **g** primary root growth for No-0 and *cepd1,2* seedlings 3 days post transfer to segmented plates with CEP3 (10^-6^ M) or water in the top segment (n = 22-23). Blue dots in **b**,**d**,**f** indicate root tip location on day of transfer to treatment plates. Letters show significant differences (ANOVA followed by Tukey HSD test, p < 0.05). Error bars show SE. Scale bars = 1 cm.

**Supplementary Table 1. Cytokinin metabolite levels in roots and shoots of wild type and *cepr1* plants**

| ***Samples*** | | ***iP*** | | | | ***iPR*** | | | | ***iPRMP*** | | | | ***iP7G*** | | | | ***iP9G*** | | | |  |  |  |  |  |  |  |  |
| --- | --- | --- | --- | --- | --- | --- | --- | --- | --- | --- | --- | --- | --- | --- | --- | --- | --- | --- | --- | --- | --- | --- | --- | --- | --- | --- | --- | --- | --- |
| Roots | No-0 | 1.78 | ± | 0.60 |  | 0.59 | ± | 0.18 |  | 2.23 | ± | 0.37 |  | 16.81 | ± | 1.06 |  | 1.93 | ± | 0.10 |  |  |  |  |  |  |  |  |  |
|  | *cepr1-1* | **3.05** | **±** | **0.81** | ***** | **1.01** | **±** | **0.29** | ***** | **2.87** | **±** | **0.39** | ***** | **14.97** | **±** | **0.82** | ***** | 2.02 | ± | 0.14 |  |  |  |  |  |  |  |  |  |
| Shoots | No-0 | 0.40 | ± | 0.05 |  | 0.61 | ± | 0.06 |  | 8.41 | ± | 0.81 |  | 36.33 | ± | 1.82 |  | 3.76 | ± | 0.23 |  |  |  |  |  |  |  |  |  |
|  | *cepr1-1* | 0.34 | ± | 0.07 |  | **0.36** | **±** | **0.07** | ******* | **5.56** | **±** | **0.73** | ******* | 34.90 | ± | 1.69 |  | 3.55 | ± | 0.22 |  |  |  |  |  |  |  |  |  |
| Roots | Col-0 | 0.89 | ± | 0.26 |  | 0.32 | ± | 0.06 |  | 0.56 | ± | 0.16 |  | 8.90 | ± | 0.45 |  | 0.83 | ± | 0.05 |  |  |  |  |  |  |  |  |  |
|  | *cepr1-3* | 0.76 | ± | 0.19 |  | **0.55** | **±** | **0.17** | ***** | **1.69** | **±** | **0.35** | ****** | 7.92 | ± | 0.78 |  | 0.75 | ± | 0.08 |  |  |  |  |  |  |  |  |  |
| Shoots | Col-0 | 0.28 | ± | 0.02 |  | 0.39 | ± | 0.11 |  | 4.63 | ± | 0.28 |  | 30.96 | ± | 2.52 |  | 3.26 | ± | 0.27 |  |  |  |  |  |  |  |  |  |
|  | *cepr1-3* | 0.24 | ± | 0.05 |  | 0.33 | ± | 0.08 |  | **3.78** | **±** | **0.44** | ***** | 27.38 | ± | 6.60 |  | 2.91 | ± | 0.16 |  |  |  |  |  |  |  |  |  |
| ***Samples*** | | ***tZ*** | | | | ***tZR*** | | | | ***tZRMP*** | | | | ***tZOG*** | | | | ***tZROG*** | | | | ***tZ7G*** | | | | ***tZ9G*** | | | |
| Roots | No-0 | 2.66 | ± | 0.55 |  | 0.30 | ± | 0.06 |  |  | *<LOD* |  |  | 1.64 | ± | 0.29 |  |  | *<LOD* |  |  | 4.53 | ± | 0.41 |  | 1.10 | ± | 0.30 |  |
|  | *cepr1-1* | **5.18** | **±** | **0.78** | ******* | **0.73** | **±** | **0.15** | ******* |  | *<LOD* |  |  | 2.04 | ± | 0.28 |  |  | *<LOD* |  |  | **5.39** | **±** | **0.52** | ***** | 1.35 | ± | 0.33 |  |
| Shoots | No-0 | 0.87 | ± | 0.18 |  | 0.30 | ± | 0.06 |  | 1.24 | ± | 0.17 |  | 1.62 | ± | 0.07 |  | 0.22 | ± | 0.05 |  | 7.10 | ± | 0.46 |  | 2.28 | ± | 0.35 |  |
|  | *cepr1-1* | 0.82 | ± | 0.24 |  | 0.25 | ± | 0.06 |  | **0.82** | **±** | **0.15** | ****** | **1.23** | **±** | **0.21** | ****** | 0.22 | ± | 0.06 |  | **5.48** | **±** | **0.64** | ****** | **1.68** | **±** | **0.20** | ***** |
| Roots | Col-0 | 0.51 | ± | 0.13 |  | 0.34 | ± | 0.07 |  | 0.95 | ± | 0.15 |  | 2.41 | ± | 0.28 |  | 0.04 | ± | 0.01 |  | 4.80 | ± | 0.28 |  | 0.90 | ± | 0.09 |  |
|  | *cepr1-3* | 0.78 | ± | 0.21 |  | **1.32** | **±** | **0.22** | ******* | **2.56** | **±** | **0.25** | ******* | **3.54** | **±** | **0.70** | ***** | 0.05 | ± | 0.01 |  | **6.62** | **±** | **1.04** | ****** | **1.56** | **±** | **0.30** | ****** |
| Shoots | Col-0 | 0.14 | ± | 0.01 |  | 0.24 | ± | 0.03 |  | 0.77 | ± | 0.12 |  | 2.13 | ± | 0.19 |  | 0.22 | ± | 0.04 |  | 7.95 | ± | 0.57 |  | 2.46 | ± | 0.24 |  |
|  | *cepr1-3* | **0.21** | **±** | **0.02** | ******* | **0.45** | **±** | **0.05** | ******* | **1.46** | **±** | **0.31** | ****** | 2.59 | ± | 0.40 |  | 0.26 | ± | 0.04 |  | **11.86** | **±** | **0.67** | ******* | **3.57** | **±** | **0.52** | ****** |
| ***Samples*** | | ***cZ*** | | | | ***cZR*** | | | | ***cZRMP*** | | | | ***cZOG*** | | | | ***cZROG*** | | | | ***cZ7G*** | | | | ***cZ9G*** | | | |
| Roots | No-0 | 1.90 | ± | 0.47 |  | 0.82 | ± | 0.24 |  | 4.88 | ± | 0.86 |  | 5.57 | ± | 0.50 |  | 1.20 | ± | 0.23 |  | 48.05 | ± | 3.76 |  | 1.09 | ± | 0.08 |  |
|  | *cepr1-1* | 2.54 | ± | 0.77 |  | 1.22 | ± | 0.35 |  | **6.53** | **±** | **0.19** | ***** | 5.76 | ± | 0.26 |  | 1.47 | ± | 0.45 |  | 43.34 | ± | 3.72 |  | **0.80** | **±** | **0.20** | ***** |
| Shoots | No-0 | 0.19 | ± | 0.02 |  | 0.36 | ± | 0.08 |  | 3.10 | ± | 0.35 |  | 1.25 | ± | 0.06 |  | 0.74 | ± | 0.06 |  | 22.14 | ± | 1.31 |  | 0.39 | ± | 0.06 |  |
|  | *cepr1-1* | 0.18 | ± | 0.03 |  | **0.21** | **±** | **0.03** | ****** | 2.81 | ± | 0.60 |  | **1.38** | **±** | **0.07** | ***** | 0.82 | ± | 0.16 |  | **26.23** | **±** | **0.93** | ******* | 0.48 | ± | 0.06 |  |
| Roots | Col-0 | 1.23 | ± | 0.40 |  | 0.51 | ± | 0.15 |  | 6.54 | ± | 0.29 |  | 2.61 | ± | 0.10 |  | 0.29 | ± | 0.04 |  | 25.20 | ± | 1.04 |  | 0.50 | ± | 0.06 |  |
|  | *cepr1-3* | 0.80 | ± | 0.17 |  | 0.40 | ± | 0.07 |  | 6.77 | ± | 2.04 |  | 2.46 | ± | 0.39 |  | 0.27 | ± | 0.03 |  | **22.68** | **±** | **1.11** | ***** | 0.51 | ± | 0.06 |  |
| Shoots | Col-0 | 0.19 | ± | 0.03 |  | 0.21 | ± | 0.05 |  | 2.58 | ± | 0.38 |  | 1.16 | ± | 0.13 |  | 1.00 | ± | 0.18 |  | 20.31 | ± | 1.75 |  | 0.34 | ± | 0.04 |  |
|  | *cepr1-3* | 0.16 | ± | 0.02 |  | 0.23 | ± | 0.04 |  | 2.04 | ± | 0.29 |  | 1.08 | ± | 0.18 |  | 1.42 | ± | 0.33 |  | 21.62 | ± | 1.89 |  | 0.40 | ± | 0.09 |  |
| ***Samples*** | | ***DHZ*** | | | | ***DHZR*** | | | | ***DHZRMP*** | | | | ***DHZOG*** | | | | ***DHZROG*** | | | | ***DHZ7G*** | | | | ***DHZ9G*** | | | |
| Roots | No-0 | 0.85 | ± | 0.24 |  | 0.21 | ± | 0.05 |  |  | *<LOD* |  |  | 0.47 | ± | 0.07 |  |  | *<LOD* |  |  |  | *<LOD* |  |  |  | *<LOD* |  |  |
|  | *cepr1-1* | 1.15 | ± | 0.34 |  | 0.29 | ± | 0.07 |  |  | *<LOD* |  |  | 0.59 | ± | 0.17 |  |  | *<LOD* |  |  |  | *<LOD* |  |  |  | *<LOD* |  |  |
| Shoots | No-0 |  | *<LOD* |  |  | 0.01 | ± | 0.00 |  |  | *<LOD* |  |  | 0.07 | ± | 0.02 |  |  | *<LOD* |  |  | 1.85 | ± | 0.25 |  |  | *<LOD* |  |  |
|  | *cepr1-1* |  | *<LOD* |  |  |  | *<LOD* |  |  |  | *<LOD* |  |  | 0.08 | ± | 0.01 |  |  | *<LOD* |  |  | 1.82 | ± | 0.10 |  |  | *<LOD* |  |  |
| Roots | Col-0 | 0.63 | ± | 0.19 |  | 0.05 | ± | 0.01 |  |  | *<LOD* |  |  | 0.49 | ± | 0.06 |  |  | *<LOD* |  |  | 1.12 | ± | 0.12 |  |  | *<LOD* |  |  |
|  | *cepr1-3* | 0.61 | ± | 0.21 |  | 0.08 | ± | 0.03 |  |  | *<LOD* |  |  | 0.52 | ± | 0.10 |  |  | *<LOD* |  |  | **1.94** | **±** | **0.30** | ******* |  | *<LOD* |  |  |
| Shoots | Col-0 | 0.21 | ± | 0.06 |  | 0.04 | ± | 0.01 |  |  | *<LOD* |  |  | 0.10 | ± | 0.03 |  |  | *<LOD* |  |  | 1.55 | ± | 0.06 |  |  | *<LOD* |  |  |
|  | *cepr1-3* | **0.67** | **±** | **0.21** | ****** | **0.02** | **±** | **0.01** | ****** |  | *<LOD* |  |  | 0.12 | ± | 0.01 |  |  | *<LOD* |  |  | **2.53** | **±** | **0.24** | ******* |  | *<LOD* |  |  |

Cytokinin content was measured in root and shoot samples from 6-d-old seedlings grown on standard ½ MS medium. Values are in pmol g^−1^ fresh weight. *<LOD*, below limit of detection. Data are mean ± S.D. (n = 5). Asterisks indicate statistically significant differences from wild type (two sample t-test; *p < 0.05, ** p < 0.01, *** p < 0.01).; *t*Z, *trans*-zeatin; iP, isopentenyladenine; *c*Z, *cis*-zeatin; DHZ, dihydrozeatin; R, riboside; RMP, riboside 5′-monophosphate; 7G, 7-glucoside; 9G, 9-glucoside; OG, *O*-glucoside.

**Supplementary Table 2. Primer sequences for qRT-PCR**

| Target | Primer sequences (F; R) |
| --- | --- |
| *EF1α* | GAGCGTTGTTCTCGTAATTGG; AAAACAGGAAGAATGTGTGTGTAGA |
| *CEPD1* | TCCGAAAAAGGGGTGGTTAT; ACTTGAACCGCATAGGACAAA |
| *CEPD2* | TCTTTGTCGGAGGCAAGC;  GCCACTAAGGTGAAGGGACA |
| *ARR5* | TCAGAGAACATCTTGCCTCG;  ATTTCACAGGCTTCAATAAGAAATC |
